# DDX3X and DDX3Y are redundant in protein synthesis

**DOI:** 10.1101/2020.09.30.319376

**Authors:** Srivats Venkataramanan, Lorenzo Calviello, Kevin Wilkins, Stephen N. Floor

## Abstract

DDX3 is a DEAD-box RNA helicase that regulates translation and is encoded by the X- and Y-linked paralogs *DDX3X* and *DDX3Y*. While DDX3X is ubiquitously expressed in human tissues and essential for viability, DDX3Y is male-specific and shows lower and more variable expression than DDX3X in somatic tissues. Heterozygous genetic lesions in DDX3X mediate a class of developmental disorders called DDX3X syndrome, while loss of DDX3Y is implicated in male infertility. One possible explanation for female-bias in DDX3X syndrome is that DDX3Y encodes a polypeptide with different biochemical activity. In this study, we use ribosome profiling and *in vitro* translation to demonstrate that the X- and Y-linked paralogs of DDX3 play functionally redundant roles in translation. We find that transcripts that are sensitive to DDX3X depletion or mutation are rescued by complementation with DDX3Y. Our data indicate that DDX3X and DDX3Y proteins can functionally complement each other in human cells. DDX3Y is not expressed in a large fraction of the central nervous system. These findings suggest that expression differences, not differences in paralog-dependent protein synthesis, underlie the sex-bias of DDX3X-associated diseases.

## Introduction

DDX3X is a ubiquitously expressed ATPase and RNA-helicase encoded by an essential gene on the X-chromosome, *DDX3X*. *DDX3X* escapes X-inactivation in a wide range of tissues (Cotton et al., 2015). Mutations in *DDX3X* are associated with numerous pathologies, including cancers like medulloblastoma (Floor et al., 2016; Kool et al., 2014; Oh et al., 2016), chronic lymphocytic leukemia (Ojha et al., 2015), squamous cell carcinoma (Stransky et al., 2011), Burkitt’s lymphoma (Gong et al., 2020), and many others (Sharma and Jankowsky, 2014). Heterozygous missense or loss-of-function mutations in *DDX3X* are also implicated in intellectual disability and autism-spectrum disorders in females (Iossifov et al., 2014; Lennox et al., 2020; Ruzzo et al., 2019; Scala et al., 2019; Snijders Blok et al., 2015; Takata et al., 2018; Wang et al., 2018; Yuen et al., 2017), with the severity of phenotype correlating with the degree of reduction in DDX3X catalytic activity (Lennox et al., 2020).

*DDX3X* has a Y-chromosome paralog, *DDX3Y*, and the protein products of these two genes are more than 90% identical (Ditton et al., 2004). *DDX3Y* is located within the *Azoospermia Factor a (AZFa)* locus nested within the non-recombining region of the Y chromosome (Kotov et al., 2017). Expression of DDX3Y alone has been shown to rescue the specific infertility phenotype caused by deletion of *AZFa* (Ramathal et al., 2015), although these findings have been disputed in mice (Zhao et al., 2019). *DDX3Y* mRNA is broadly expressed across tissues, but activation of a testis-specific distal promoter produces a transcript isoform with a distinct 5′ leader that also contributes to its translational control (Jaroszynski et al., 2011). Because of this, expression of DDX3Y was thought to be testis-specific, but later work suggested that DDX3Y is expressed more broadly in tissues across the human body (Uhlen et al., 2015)(data available from https://v19.proteinatlas.org). Furthermore, recent evidence suggests that depletion of DDX3Y results in neural differentiation defects, indicating wider tissue distribution of DDX3Y than previously assumed (Vakilian et al., 2015). Notably, *DDX3Y* expression is consistently lower than that of its X-linked paralog (Uhlen et al., 2015)(data available from https://v19.proteinatlas.org).

*DDX3X* and *DDX3Y* both encode for variants of DDX3, a DEAD-box RNA chaperone that facilitates translation initiation on mRNAs with structured 5′ leaders (Calviello et al., 2020; Oh et al., 2016). We recently defined the set of genes that depend on DDX3X for their translation and proposed a model where this translation is facilitated by the resolution of these highly structured 5′ leaders by 40S-associated DDX3X (Calviello et al., 2020), consistent with prior work (Chen et al., 2018). However, the role of DDX3Y in translation and gene expression – and how similar or different its function is to DDX3X - remains incompletely understood.

Genetic diseases involving *DDX3X* such as DDX3X syndrome demonstrate severe sex bias (n=107, 104 females, 3 males). This cohort contains *de novo* genetic lesions in *DDX3X* that are exclusively heterozygous (or hemizygous in males) (Lennox et al., 2020). These observations have been attributed to the inability of the Y-linked DDX3 paralog to functionally complement DDX3X, leading to embryonic lethality in males with inactivating mutations of DDX3X – whereas heterozygous females are viable, but suffer from symptoms of varying intensity. *Ddx3y* is unable to compensate for the loss of *Ddx3x* during embryonic development in mice (Chen et al., 2016) This is in contrast to the observation that mouse *Ddx3y i*s able to compensate for the loss of *Ddx3x* during neural development, insulating male mice from ataxia and seizure phenotypes upon ablation of *Ddx3x*, but is unable to reverse the susceptibility to hindbrain malignancies conferred by *Ddx3x* loss (Patmore et al., 2020). Conversely, the loss of DDX3Y confers infertility in human males, despite the robust expression of DDX3X in germ-line tissues (Ramathal et al., 2015). In principle, sex bias in DDX3X-dependent disorders could arise from differential expression between DDX3X and DDX3Y across tissues, different activities of their gene products, dosage compensation by the wild-type locus, or other unknown mechanisms.

In this study, we tested the functions of DDX3X and DDX3Y proteins in translation through cell based and biochemical assays. We depleted endogenous DDX3X in male HCT 116 cells using an inducible degron system, complemented with either *DDX3X* or *DDX3Y* cDNAs, and measured translation and RNA abundance using ribosome profiling and RNA-seq. We find that the substitution of DDX3X with DDX3Y does not significantly affect translation or transcript levels. Using an in vitro reporter system, we demonstrate that genes that are robustly susceptible to depletion of DDX3 do not show significant changes upon substitution of DDX3X with DDX3Y. Taken together with their distinct tissue-level expression patterns, our data suggests that the X- and Y-linked paralogs of DDX3 are redundant in protein synthesis, implying tissue-specific variation of DDX3Y expression or another mechanism beyond translational control underlies the sex bias of DDX3X associated developmental disorders.

## Materials and methods

### DDX3 sequence alignment and phylogenetic tree

Reference protein sequences of human DDX3X and DDX3Y were obtained from RefSeq and alignment was performed using the MAFFT algorithm using the BLOSUM62 scoring matrix with default parameters (gap open penalty = 1.53; gap extension penalty = 0.123) (Katoh et al., 2002). Protein sequences for mammalian DDX3 orthologs (both X and Y, where available and reliably annotated) were obtained using a combination of the NCBI Gene ortholog finder and PSI-blast (Altschul et al., 1997). Multiple sequence alignment was performed as described above, and a phylogenetic tree was constructed using the Interactive Tree of Life (iTOL) tool (v5.6.3) (Letunic and Bork, 2019).

### Construction of DDX3X-degron cell line

HCT-116 cells expressing OsTIR1 were constructed as described in (Natsume et al., 2016). Briefly, a plasmid co-expressing gRNA for the AAVS1 safe-harbor locus and SpCas9 was co-transfected with plasmid encoding for OsTIR1 and homologous arms for the HDR at the AAVS1 locus. Selection was performed in 1 μg/ml of puromycin, colonies were picked and validated by PCR and western blot.

The HCT-116 OsTIR1 expressing cell line was transfected with 800 ng (total) of an equimolar mixture of pX330-based CRISPR/Cas plasmid with guide RNAs targeting the C-terminal end of DDX3X and 1200 ng of a short homology donor carrying the mAID degron tag with a G418 resistance cassette in a pUC19 backbone. Selection was performed in 700 μg/ml of G418, and single colonies picked and validated by PCR and western blot. Selection, picking, clonal expansion and validation were repeated twice (total of 3 rounds) to ensure a clonal cell line.

### Induction of DDX3X degradation and transfection of DDX3X-FLAG or DDX3Y-FLAG

Partially codon-optimized DDX3X-FLAG and DDX3Y-FLAG were obtained from Twist Biosciences and cloned into an expression vector under the control of a CMV promoter. For ribosome profiling and *in vitro* translation lysates, expression plasmids (50μg of DDX3X-FLAG and 75μg of DDX3Y-FLAG per 15 cm plate, to equalize expression levels) were transfected into cells using a 3:1 ratio of lipofectamine 2000 to plasmid. 24 hours post-transfection, 500 μM Indole-3-acetic acid (IAA, Grainger, 31FY95) was added to the cells (in fresh media) from a 500 mM stock in DMSO. Protein levels were assayed by western blot.

### Antibodies

Primary antibodies used in this study include rabbit polyclonal anti-DDX3 (1:5000 in 5% milk, custom made by Genemed Synthesis using peptide ENALGLDQQFAGLDLNSSDNQS) (Lee et al., 2008), anti-actin HRP (1:10000 in 5% milk, Santa Cruz Biotechnology, sc-47778), anti-FLAG HRP (1:10000 in 5% milk, Sigma, A8592).

### Ribosome profiling library construction

DDX3X-mAID tagged HCT-116 (one 15 cm dish at 80-90% confluency per replicate) cells expressing OsTIR1 were transfected with either DDX3X or DDX3Y. 24 hours post-transfection, media was changed and fresh media with 500 μM IAA was added to cells. Un-transfected cells were treated with either DMSO or IAA. 48 hours after auxin addition, cells were treated with 100 μg/ml cycloheximide (CHX) and harvested and lysed as described in (McGlincy and Ingolia, 2017). Briefly, cells were washed with PBS containing 100 μg/ml CHX and lysed in ice-cold lysis buffer (20 mM TRIS-HCl pH 7.4, 150mM NaCl, 5 mM MgCl2, 1mM DTT, 100 μg/ml CHX, 1 % (v/v) Triton X-100, 25 U/ml TurboDNase (Ambion)). 240 μl lysate was treated with 6 μl RNase I (Ambion, 100 U/μl) for 45 minutes at RT with gentle agitation and further digestion halted by addition of SUPERase:In (Ambion). Illustra Microspin Columns S-400 HR (GE healthcare) were used to enrich for monosomes, and RNA was extracted from the flow-through using Direct-zol kit (Zymo Research). Gel slices of nucleic acids between 24-32 nts long were excised from a 15% urea-PAGE gel. Eluted RNA was treated with T4 PNK and preadenylated linker was ligated to the 3’ end using T4 RNA Ligase 2 truncated KQ (NEB, M0373L). Linker-ligated footprints were reverse transcribed using Superscript III (Invitrogen) and gel-purified RT products circularized using CircLigase II (Lucigen, CL4115K). rRNA depletion was performed using biotinylated oligos as described in (Ingolia et al., 2012) and libraries constructed using a different reverse indexing primer for each sample.

RNA was extracted from 25 μl intact lysate (non-digested) using the Direct-zol kit (Zymo Research) and stranded total RNA libraries were prepared using the TruSeq Stranded Total RNA Human/Mouse/Rat kit (Illumina), following manufacturer’s instructions.

Libraries were quantified and checked for quality using a Qubit fluorimeter and Bioanalyzer (Agilent) and sequenced on a HiSeq 4000 sequencing system (single end, 65 nt reads).

### Pre-processing and alignment of NGS data

All NGS data was processed as previously described (Calviello et al., 2020). Briefly, adapter sequences were trimmed from the footprint reads, UMI sequences were removed and reads were collapsed. Reads aligned to rRNA or a collection of snoRNAs, tRNAs and miRNAs were discarded. The filtered reads were mapped to the hg38 version of the human genome using STAR (Dobin et al., 2013) and count matrices built as described previously (Calviello et al., 2020) using Ribo-seQC (Calviello et al., 2019). Only reads above 24 nucleotides long were included in the count. Read counts for all libraries are in Table S1.

### Differential expression analysis

Differentially expressed genes between the samples were identified using DESeq2 (Love et al., 2014). Only genes with average normalized number of counts greater than 20 for both RNA and Ribo-seq were retained. Genes with transcript levels were defined as either “RNA_up” or RNA_down”, with an adjusted p-value cutoff of 0.01. Differential translation regulation was calculated using DESeq2 interaction terms and a likelihood ratio test, with an adjusted p-value cutoff of 0.05 and differentially translated genes were defined as “TE-up” or TE_down”. DE details for all genes that met the read count cutoff are in Table S2.

### Transcript features analysis

5′UTR ribosome skew (riboskew) was calculated as previously defined (Calviello et al., 2020). Effect sizes (non-parametric Cliff’s Delta measure) between the transcript classes were calculated using the ‘effsize’ R package (Torchiano, 2020).

### In vitro transcription, capping, and 2’-O methylation of reporter RNAs

Annotated 5′ UTRs for selected transcripts were cloned upstream of Renilla Luciferase (RLuc) under the control of a T7 promoter, with 60 adenosine nucleotides downstream of the stop codon to mimic polyadenylation. Untranslated regions were cloned using synthetic DNA (Integrated DNA Technologies) or by isolation using 5′ RACE (RLM-RACE kit, Invitrogen). Template was PCR amplified using Phusion polymerase from the plasmids using the following primers, and gel purified, as described (Floor and Doudna, 2016).

pA60 txn rev: TTT TTT TTT TTT TTT TTT TTT TTT TTT TTT TTT TTT TTT TTT TTT TTT TTT TTT TTT TTT CTG CAG

pA60 txn fwd: CGG CCA GTG AAT TCG AGC TCT AAT ACG ACT CAC TAT AGG

100 uL in vitro transcription reactions were set up at room temperature with 1-5 micrograms of purified template, 7.5mM ACGU ribonucleotides, 30mM Tris-Cl pH 8.1, 125mM MgCl2, 0.01% Triton X-100, 2mM spermidine, 110mM DTT, T7 polymerase and 0.2 U/uL units of Superase-In RNase inhibitor (Thermo-Fisher Scientific). Transcription reactions were incubated in a PCR block at 37 degrees C for 1 hour. 1 uL of 1 mg/mL pyrophosphatase (Roche) was added to each reaction, and the reactions were subsequently incubated in a PCR block at 37 degrees C for 3 hours. 1 unit of RQ1 RNase-free DNase (Promega) was added to each reaction followed by further incubation for 30 minutes. RNA was precipitated by the addition of 200 uL 0.3M NaOAc pH 5.3, 15 ug GlycoBlue co-precipitant (Thermo-Fisher Scientific) and 750 uL 100% EtOH. Precipitated RNA was further purified over the RNA Clean & Concentrator-25 columns (Zymo Research). Glyoxal gel was run to assess the integrity of the RNA before subsequent capping and 2’ O-methylation.

20 ug of total RNA was used in a 40 uL capping reaction with 0.5mM GTP, 0.2mM s-adenosylmethionine (SAM), 20 units of Vaccinia capping enzyme (New England Biolabs), 100 units of 2’-O-Me-transferase (New England Biolabs) and 25 units RNasin Plus RNase inhibitor (Promega). The reactions were incubated at 37 degrees C for 1 hour, followed by purification over the RNA Clean & Concentrator-25 columns (Zymo Research) and elution in DEPC H2O. Glyoxal gel was run to assess the integrity of the RNA before proceeding to in vitro translation reactions.

### Preparation of cellular extracts for in vitro translation

Three 150mm plates of HCT 116 cells were trypsinized and pelleted at 1000g, 4 degrees C. One cell-pellet volume of lysis buffer (10mM HEPES, pH 7.5, 10mM KOAc, 0.5mM MgOAc2, 5mM DTT, and 1 tablet miniComplete EDTA free protease inhibitor (Roche) per 10 mL) was added to the cell pellet and was incubated on ice for 45 minutes. The pellet was homogenized by trituration through a 26G needle attached to a 1 mL syringe 13-15 times. Efficiency of disruption was checked by trypan blue staining (>95% disruption target). The lysate was cleared by centrifugation at 14000g for 1 minute at 4 degrees C, 2-5 ul was reserved for western blot analysis, and the remainder was aliquoted and flash frozen in liquid nitrogen. Preparation of HEK293T lysates with DDX3X siRNA knockdown and mutant expression has been previously described (Lennox et al., 2020)

### In vitro translation

5 uL in vitro translation reactions were set up with 2.5 uL of lysate and 20 ng total RNA (0.84mM ATP, 0.21mM GTP, 21mM Creatine Phosphate, 0.009units/mL Creatine phosphokinase, 10mM HEPES pH 7.5, 2mM DTT, 2mM MgOAc, 100mM KOAc, 0.008mM amino acids, 0.25mM spermidine, 5 units RNasin Plus RNase inhibitor (Promega) as described (Lee et al., 2015). Reaction tubes were incubated at 30 degrees C for 45 minutes, and expression of the reporter was measured using the Renilla Luciferase Assay System (Promega) on a GloMax Explorer plate reader (Promega). Drug treatments were performed by incubating the reaction with the indicated compound for 15 minutes at 30 degrees C prior to the addition of RNA.

### Analysis of FANTOM5 tissue level expression data

FANTOM5 per gene transcript expression levels (CAGE data) was obtained based on The Human Protein Atlas version 19.3 and Ensembl version 92.38 (Lizio et al., 2015). Quantile-normalized and scaled TPMs were plotted for common tissues as well as male and female specific tissues.

## Results

The DEAD-box RNA helicase DDX3 has two paralogs, one each on each of the sex chromosomes. The X-linked *DDX3X* gene is located on a non-pseudoautosomal region of the short arm of the X-chromosome (**Figure 1A**). *DDX3Y*, a paralog of *DDX3X*, is located on the long arm of the Y chromosome in the male-specific region (**Figure 1A**). The sex bias of DDX3X syndrome suggests that males with inactivating mutations in DDX3X suffer embryonic lethality, implying that human *DDX3Y* cannot functionally complement *DDX3X*. However, *DDX3X* and *DDX3Y* protein products show ~92% homology, with most of the differences in the N-terminal domain (**Figure 1B**), which is dispensable for RNA duplex unwinding (Floor et al., 2016). Nevertheless, a phylogenetic tree of available mammalian DDX3 protein sequences reveals significant homology between mammalian DDX3Y, and separation of the “DDX3Y clade” away from DDX3X (**Figure 1C, Figure S1**). In many cases, DDX3Y sequences of evolutionary distant mammals show greater homology than the paralogs within a given species (**Figure 1C, Figure S1**). This indicates both a high degree conservation of the DDX3 orthologs as well as a potential function for DDX3Y distinct from its X-linked paralog.

**Figure 1:**
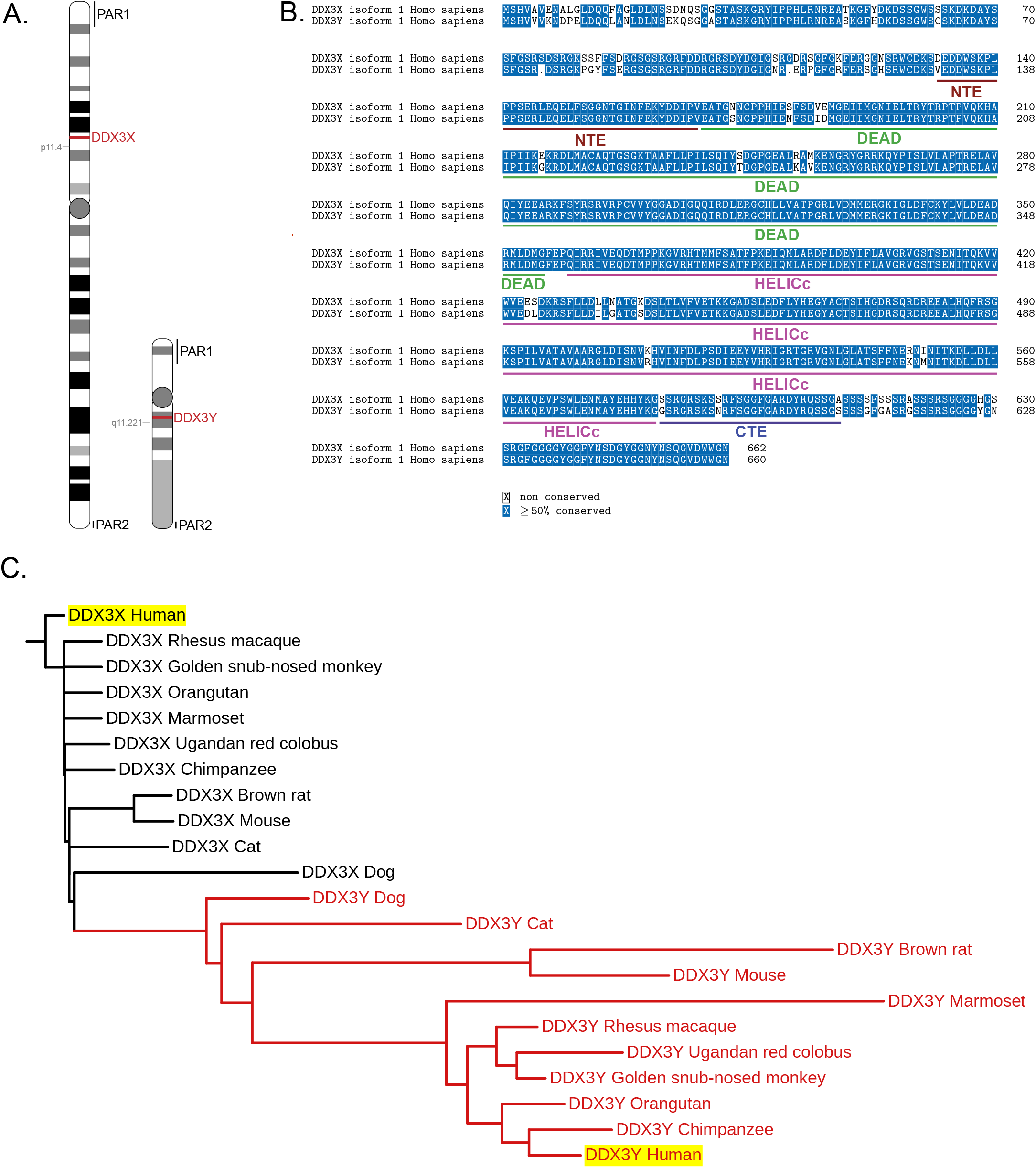
**A**. Graphic indicating the locations of *DDX3X* and *DDX3Y* on their respective chromosomes. Pseudoautosomal regions (PAR) are indicated **B**. Alignment of human DDX3X and DDX3Y demonstrating ~92% sequence identity. Domain architecture of DDX3X is indicated **C**. Phylogenetic tree indicating distances between the sequences of DDX3X and DDX3Y in mammals (where sequence is available, only selected species indicated, see Figure S1 for tree of all available mammalian sequences). The cluster of mammalian DDX3Y orthologs is indicated with red branches. Human orthologs are highlighted in yellow.

In order to dissect the similarities and differences between the functions of DDX3X and DDX3Y, we used a cell line we generated to rapidly and efficiently degrade endogenous DDX3 protein X in human male-derived colorectal cancer HCT 116 cells, which show a stable diploid karyotype (Natsume et al., 2016; Waldman et al., 1995). We tagged DDX3X with a 68 amino acid fragment of the Auxin Inducible Degron (AID) tag termed mini-AID (mAID) in parental cells expressing the auxin-responsive F-box protein, TIR1, from *Oryza sativa* (OsTIR1) (Natsume et al., 2016). Cells treated with the synthetic auxin, Indole-3-acetic acid (IAA) exhibit >90% reduction in DDX3X protein levels by 12 hours of treatment (Calviello et al., 2020) (**Figure 2A**). Using this system, we have previously shown that depletion of DDX3X (48-hour auxin treatment only) affects the translation of a subset of cellular messages (Calviello et al., 2020) (**Figure S2A**). Transcripts with increased DDX3X sensitivity have low basal translation efficiency and increased GC content and RNA structure within their 5′ leader sequences (Calviello et al., 2020).

**Figure 2:**
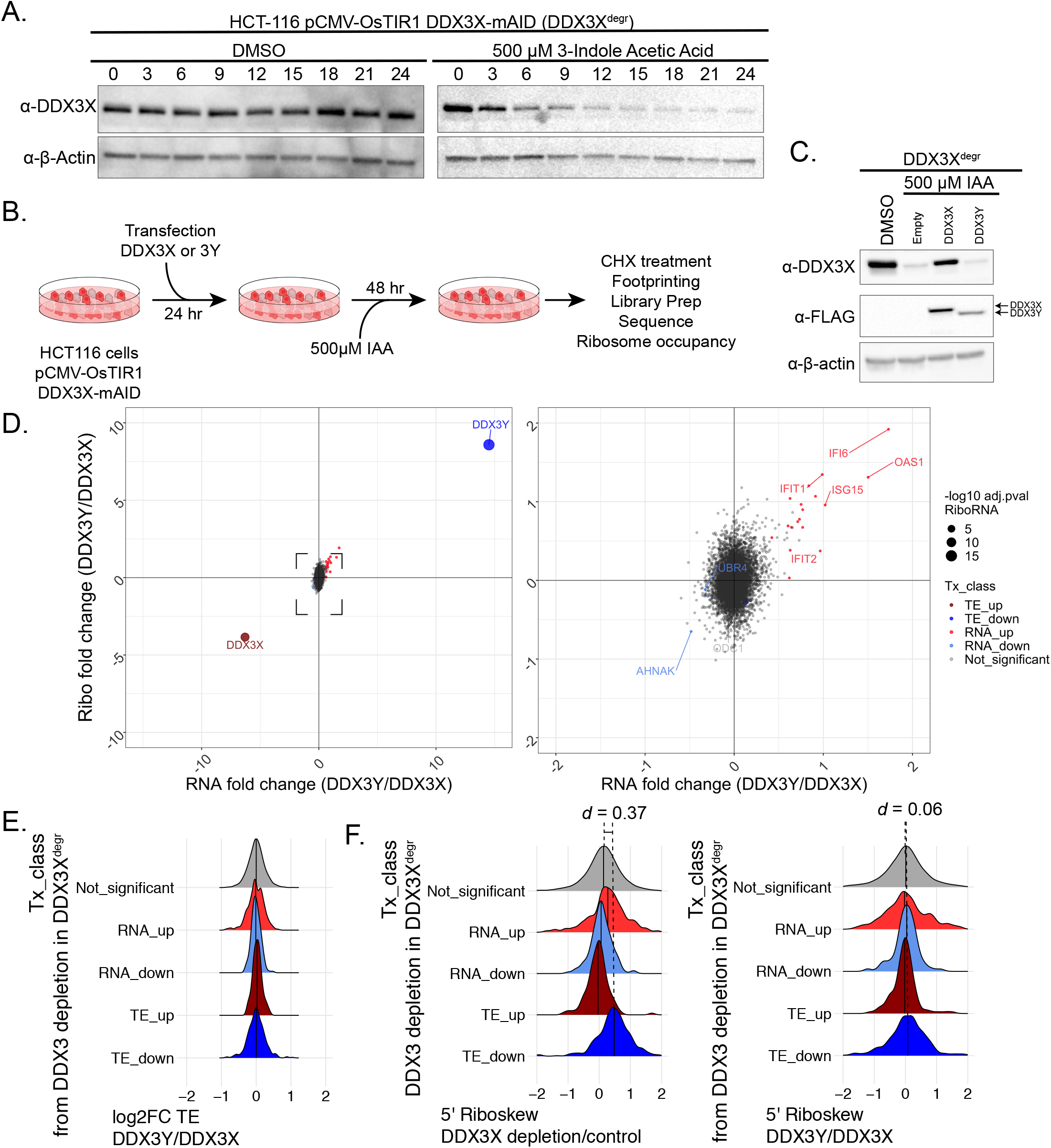
**A**. Western blot for degradation of endogenous DDX3X tagged with mini-AID (mAID) upon treatment with an auxin (indole-3-acetic acid) in HCT-116 human colorectal carcinoma cell line expressing OsTIR1 under the control of a CMV promoter (DDX3^degr^). **B**. Experimental schematic for ribosomal profiling after replacement of endogenous DDX3X with exogenous DDX3X or 3Y. **C**.. Western blot for expression of exogenous FLAG-tagged DDX3X and DDX3Y in HCT-116 cells after degradation of endogenous DDX3X. **D. Left:** Full and **Right:** zoomed in plots of differential expression analysis of RNA and ribosome profiling changes upon complementation of DDX3X with either DDX3Y or DDX3X. Point size indicates p-value of differential translation. **E**. Fold-change in TE between DDX3Y and DDX3X expression in mRNAs classified based on DDX3-sensitivity (as determined by response in DDX3X depletion). All transcript classes show no change in TE between DDX3Y and DDX3X expression. **F**. The fold-change of the ratio in ribosome occupancy in the 5’ UTR versus the coding sequence **Left:** DDX3-sensitive mRNAs (TE_down in DDX3X depletion) have increased ratio of ribosomes in the 5’ UTR compared to the CDS. **Right:** The same mRNAs show no change in the ribosome occupancy ratio upon expression of DDX3Y compared to DDX3X. Effect size (Cliff’s delta) between the “Not_significant” and “TE_down” groups is indicated.

Transient transfections with either FLAG-tagged DDX3X or DDX3Y cDNAs followed by degradation of endogenous DDX3X protein (48-hour auxin treatment beginning 24 hours post-transfection) allowed for almost complete replacement of the endogenous DDX3 protein with exogenous DDX3X or DDX3Y (**Figure 2B-C**). We measured transcript levels and ribosome occupancy under these experimental conditions using ribosome profiling and RNA-seq. Expression of DDX3Y shows little to no changes in ribosome occupancy across the entire transcriptome when compared to DDX3X (**Figure 2D**). Upregulated genes in DDX3Y vs DDX3X cells are related to the increased amount of DDX3Y-expressing plasmid that was required to be transfected to equalize protein levels (see Materials and Methods) (**Figure S2B-C**).

To increase statistical power to detect a difference between DDX3X and DDX3Y, we tested for differences between groups of transcripts. DDX3X sensitive transcripts show no change in their translation efficiency upon substitution of DDX3X with DDX3Y (**Figure 2E**; TE_down in depletion vs. control without transfection; Calviello et al., 2020). Additionally, DDX3X-sensitive transcripts show increased ribosome occupancy in 5′ leaders compared to the CDS, consistent with previous observations (Calviello et al., 2020). However, DDX3X-sensitive transcripts show no change in the ratio of ribosomes in the 5′ leader of the transcript to the coding sequence (5′ riboskew) upon the expression of DDX3Y compared to DDX3X (**Figure 2F**). Furthermore, transcripts with the largest magnitude of raw changes in TE between DDX3Y and DDX3X expression (without considering statistical significance) show no propensity for increased 5′ UTR GC content, a signature of DDX3 sensitivity, consistent with the marginal changes being due to a non-specific effect (**Figure S2D-E**). These observations suggest that the X- and Y-linked paralogs of DDX3 perform redundant functions in translation.

In order to further test the impact of DDX3X and DDX3Y on translation, we cloned a set of DDX3-sensitive 5′ UTRs (defined in (Calviello et al., 2020) upstream of a Renilla luciferase reporter and performed in vitro transcription, capping and 2′-O methylation to generate a set of reporter RNAs. We made translation extracts from HCT 116 cells either depleted of endogenous DDX3X or depleted and complemented by transient transfection of exogenous DDX3X and DDX3Y as described previously (**Figure 3A-B**; Calviello et al., 2020). As previously observed, 5′ UTRs of DDX3 sensitive mRNAs also confer DDX3 dependence to a luciferase reporter, while the control RNA with a simple 5′ UTR remains unaffected (**Figure 3C**). Consistent with the ribosome profiling results, DDX3 sensitive 5′ UTRs show no change in reporter output when translated with extracts containing only DDX3Y compared to DDX3X (**Figure 3D**). This includes the ODC1 reporter, which is sensitive to every perturbation in DDX3X that we have tested, included depletion and inactivating mutants (Calviello et al., 2020); **Figure 3E**). These data further reinforce the notion that DDX3X and DDX3Y are functionally redundant in regulation of translation.

**Figure 3:**
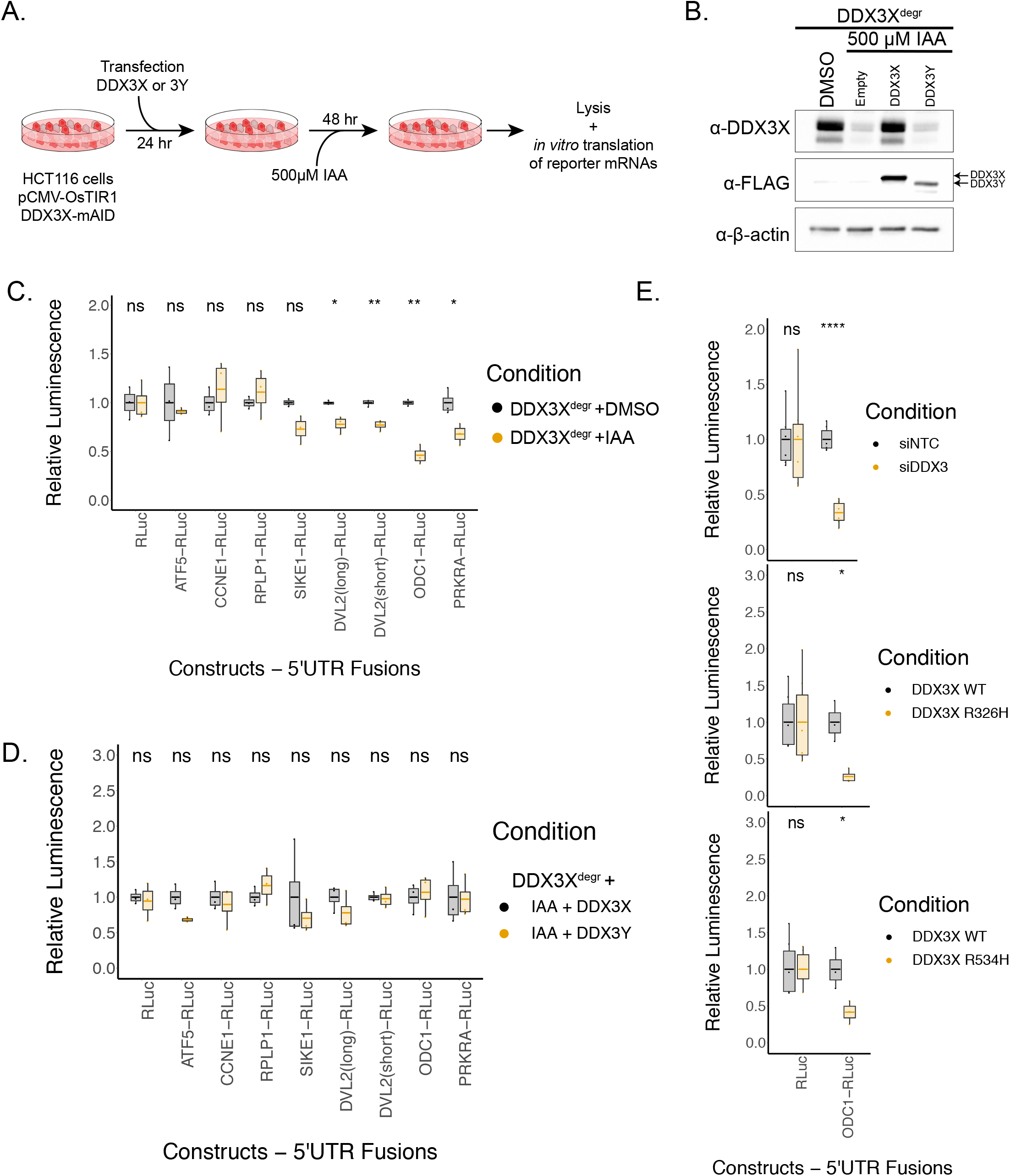
**A**. Experimental schematic for *in vitro* translation after replacement of endogenous DDX3X with exogenous DDX3X or 3Y. **B**. Western blot for degradation of endogenous DDX3X tagged with mini-AID (mAID) and expression of exogenous FLAG-tagged DDX3X and DDX3Y in HCT-116 cells after degradation of endogenous DDX3X in *in vitro* translation extracts. **C**. Translation of in vitro transcribed reporter RNAs in control (DMSO) or DDX3X-degraded (auxin treated) HCT-116 cells. **D**. Translation of in vitro transcribed reporter RNAs in lysates from either FLAG-tagged DDX3X or DDX3Y on top of depletion of endogenous DDX3X. Translation of DDX3X sensitive RNAs show no change in reporter output in the presence of DDX3Y vs. DDX3X. **E**. Translation of in vitro transcribed Renilla luciferase reporter RNA fused to the ODC1 5’ leader in lysates derived from HEK293T cells treated with and siRNA against DDX3X vs. control (top), or various inactivating DDX3X mutants compared to WT.

The results above suggest that DDX3Y can rescue translational deficiencies caused by loss of DDX3X. To reconcile this with the observed sex-bias in DDX3X-dependent disorders, we examined FANTOM-CAGE expression data for the paralogs across multiple human tissues (Lizio et al., 2015). Our examination confirmed that as previously reported, expression of *DDX3Y* is prominent in the tissues of the male reproductive system (prostate, seminal vesicle, testes etc.; **Figure 4A, Figure S3**). Furthermore, *DDX3Y* expression is undetectable in several cell types in the brain, including the midbrain, hippocampus, thalamus and amygdala (**Figure 4A, Figure S3**). Taken together, we found that *DDX3X* and *DDX3Y* functioned similarly in translation and are differentially expressed at the RNA level between human tissues.

**Figure 4:**
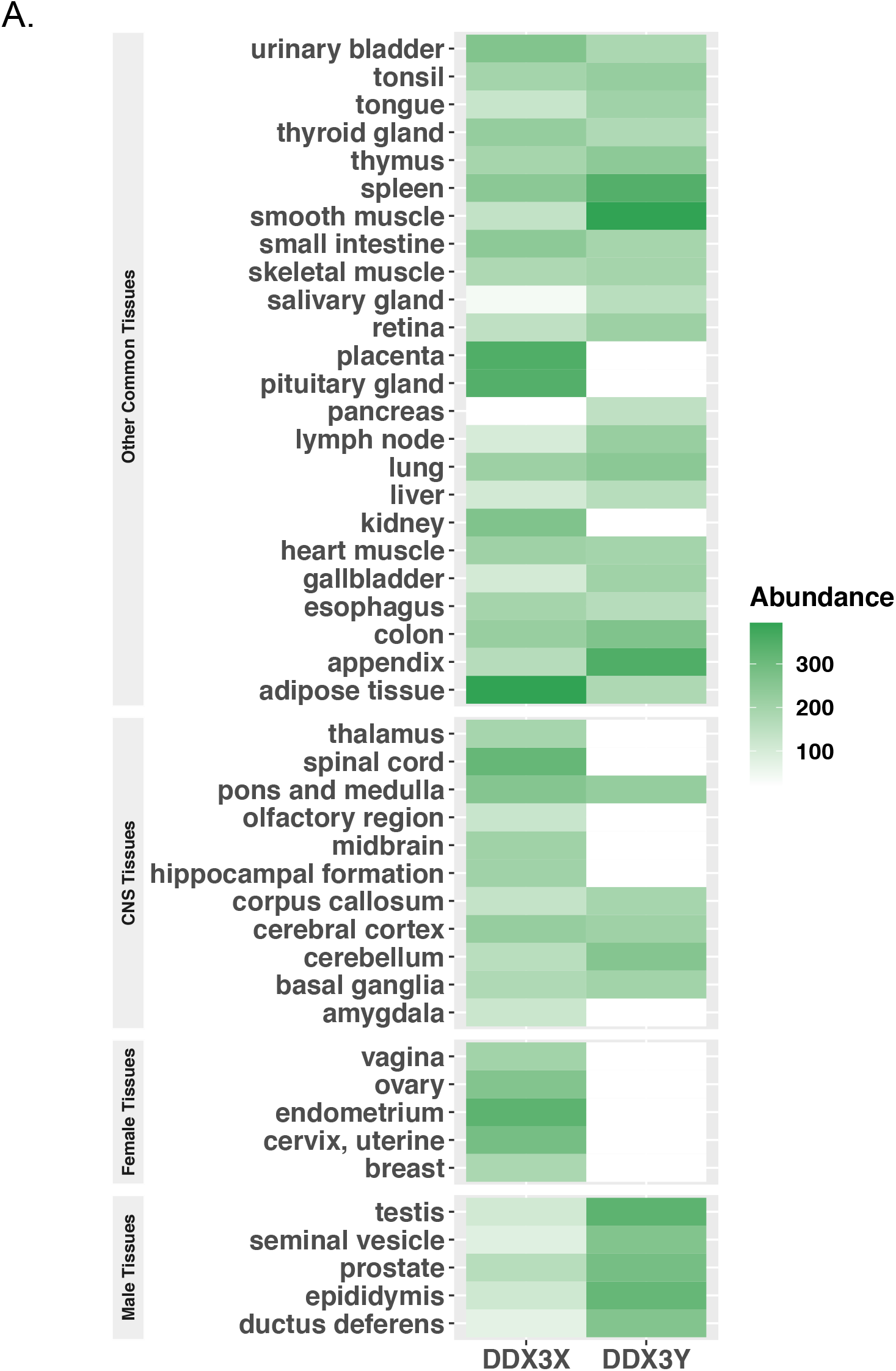
**A**. Quantile-normalized expression of DDX3X and DDX3Y across a number of human tissues (data from GTeX). DDX3Y expresson is particularly prominent in male-specific tissue types such as prostate and seminal vesicles, in the testes, and notably deplete in numerous tissues of the central nervous system.

## Discussion

The DEAD-box protein DDX3 is encoded by two paralogous genes in humans, *DDX3X* and *DDX3Y,* the protein products of which show 92% sequence identity (**Figure 1B**). Heterozygous inactivating mutations in *DDX3X* are linked to DDX3X syndrome, a neurodevelopmental delay and autism-spectrum disorder (Lennox et al., 2020). DDX3X syndrome shows a severe sex bias with >95% of known cases affecting females. This sex bias coupled with the observation that inactivating mutations are heterozygous in females (Lennox et al., 2020), suggests that *DDX3Y* is unable to compensate for the impairment of DDX3X function in brain despite recent observations that DDX3Y may also modulate neural development (Vakilian et al., 2015). In fact, the presumed causative *DDX3X* mutations in males diagnosed with DDX3X syndrome shows only very mild impairment of protein function, such that the heterozygous mother remains asymptomatic (Kellaris et al., 2018; Snijders Blok et al., 2015). This further indicates that *DDX3Y* can only partially complement mutations in *DDX3X* function in the brain. Conversely, loss of *DDX3Y* results in a male infertility phenotype despite robust expression of DDX3X in male germline tissues, indicating that *DDX3X* cannot complement loss of *DDX3Y* in the male reproductive system (Kotov et al., 2017; Ramathal et al., 2015; Rauschendorf et al., 2014)

Using both siRNA depletion and an auxin-inducible degron, we have previously demonstrated that DDX3X is required for translation initiation on transcripts with structured 5′ leader sequences (Calviello et al., 2020). In this study, we examined the global translation profiles when cells depleted of endogenous DDX3X are complemented with cDNAs of either DDX3X or DDX3Y. We could not detect significant changes of translation or steady state RNA levels (**Figure 2**). We used *in vitro* translation experiments with a luciferase reporter to confirm 5′ leaders of genes that are sensitive to either depletion of DDX3X or inactivating mutations show no significant differences in translation in extracts containing either DDX3X or DDX3Y (**Figure 3, Figure S2–S3**). In fact, the genes that show decreased translation (TE_down) upon depletion of DDX3X show an increased change in the relative ribosome occupancy in the 5′ leader compared to the CDS, a signature that is lost while comparing DDX3Y to DDX3X (**Figure 2**). This suggests that the two human paralogs of DDX3 perform equivalent functions in translation, at least in the context of our experimental system.

At least four models could potentially explain the inability of the DDX3 paralogs to complement each other. First, the two paralogs of DDX3 could have distinct target complements and affect the translation of different sets of transcripts. Second, the two paralogs could have the same target propensity but distinct subcellular localizations, and thus differ in their effects. These possibilities are partially rebuffed by the gain of DDX3Y expression complementing for loss of DDX3X in certain lymphoid malignancies (Gong et al., 2020) and in BHK21 hamster cells (Sekiguchi et al., 2004). Additionally, a genome-wide CRISPR screen for essential genes identified DDX3Y as being essential only in Raji cells (which are also of hematopoietic origin), which had suffered a truncating mutation in DDX3X, further indicating genetic complementation (Wang et al., 2015). However, genetic complementation might mask a subtle fitness defect or gene expression changes that could be important in other cellular states. In this study, we quantitatively demonstrate that the translation profile of DDX3X and DDX3Y-expressing cells are functionally identical, arguing against the two scenarios laid out above.

A third model to explain the lack of paralog complementation at the organism level is tissue specific expression patterns of the paralogs. At the RNA level, *DDX3Y* expression is sporadic in the different cell types of the brain, as well as all tissues excepting those of the male reproductive system (**Figure 4**). It has previously been suggested that specific transcriptional and translational regulation contribute to increased DDX3Y protein levels in the testes (Jaroszynski et al., 2011; Rauschendorf et al., 2011). One potential explanation for the inability of *DDX3Y* to complement *DDX3X* in the brain could be that the protein is expressed in insufficient quantities. Conversely, although DDX3X is detectable in adult male testes, multiple layers of transcriptional and translational control restrict its expression to post-meiotic spermatids (Rauschendorf et al., 2014). The absence of DDX3X protein in premeiotic spermatids could explain its inability to rescue the infertility phenotype conferred by *DDX3Y* loss. A recent analysis of tissue specific expression of sex-linked paralogs correcting for multi-mapping short sequencing reads estimated that while the cumulative expression of DDX3 is statistically identical across tissues for individuals with XX and XY genotypes, the DDX3Y expression makes up only a small minority of total DDX3 (Godfrey et al., 2020). This lends further support to the idea that while the two paralogs are functionally identical, spatiotemporal differences in their expression precludes complementation in a number of scenarios. It is important to note that this model requires future work to be confirmed. Lineage-specific complementation of DDX3 paralogs *in vivo* would formally test this hypothesis, and there are already hints of paralog-specific developmental effects (Patmore et al., 2020).

A fourth model that cannot be formally ruled out is that there are specific context-dependent factors in neurodevelopment and germ-cell development that cause differential translational regulation by the X- and Y-linked DDX3 paralogs, and that these contribute to their inability to complement each other in these contexts – despite the gene products of *DDX3X* and *DDX3Y* having redundant functions. Some evidence in support of this model comes from the observation that in mice, while *Ddx3y* is able to compensate for the neurodevelopmental phenotypes caused by the loss of *Ddx3x* in neural lineages, it is unable to suppress the increase in hindbrain malignancies also caused by *Ddx3x* loss (Patmore et al., 2020). This would indicate some form of context-dependent accessory factor influencing the target complement of Ddx3 homologs. However, it is worth noting that the ability of male mice with genetic lesions in *Ddx3x* to survive without neurological anomalies has not been recapitulated in humans as genetic variation in *DDX3X* in the human population is extremely low (Karczewski et al., 2020). We suggest that some combination of tissue-specific paralog and unknown co-factor expression may explain the sex bias of DDX3X associated developmental disorders, but that inherent differences in molecular function are unlikely. Further investigations in these directions could reveal specific targets or factors that could be exploited to mitigate the phenotypic consequences of genetic lesions in the *DDX3* paralogs.

## Data access

Sequencing data can be retrieved using GEO accession number GSE180669. Source code for the analysis of ribosome profiling data was adapted from: https://github.com/lcalviell/DDX3X_RPCLIP. Source code specific to the analysis in this manuscript is available at: https://github.com/srivats-venkat/DDX3X_3Y_comparison.

## Author contributions

Conceptualization, S.V. and S.N.F.; Investigation, S.V. and L.C.; Writing – Original Draft, S.V.; Writing – Review & Editing, S.N.F., S.V. and L.C.; Resources, S.V. and K.W.; Funding Acquisition, S.N.F.; Supervision, S.N.F.

## Acknowledgements

We thank members of the Floor lab for feedback on the manuscript. Computation was supported by the UCSF Wynton computing infrastructure. This work was supported by the UCSF Program for Breakthrough Biomedical Research, funded in part by the Sandler Foundation (to SNF), the California Tobacco-Related Disease Research Grants Program 27KT-0003 (to SNF), the National Institutes of Health DP2GM132932 (to SNF), the National Institutes of Health F32GM133144 (to SV), and the California Tobacco-Related Disease Research Grants Program T30DT1004 (to KW).

---

**Figure S1:**
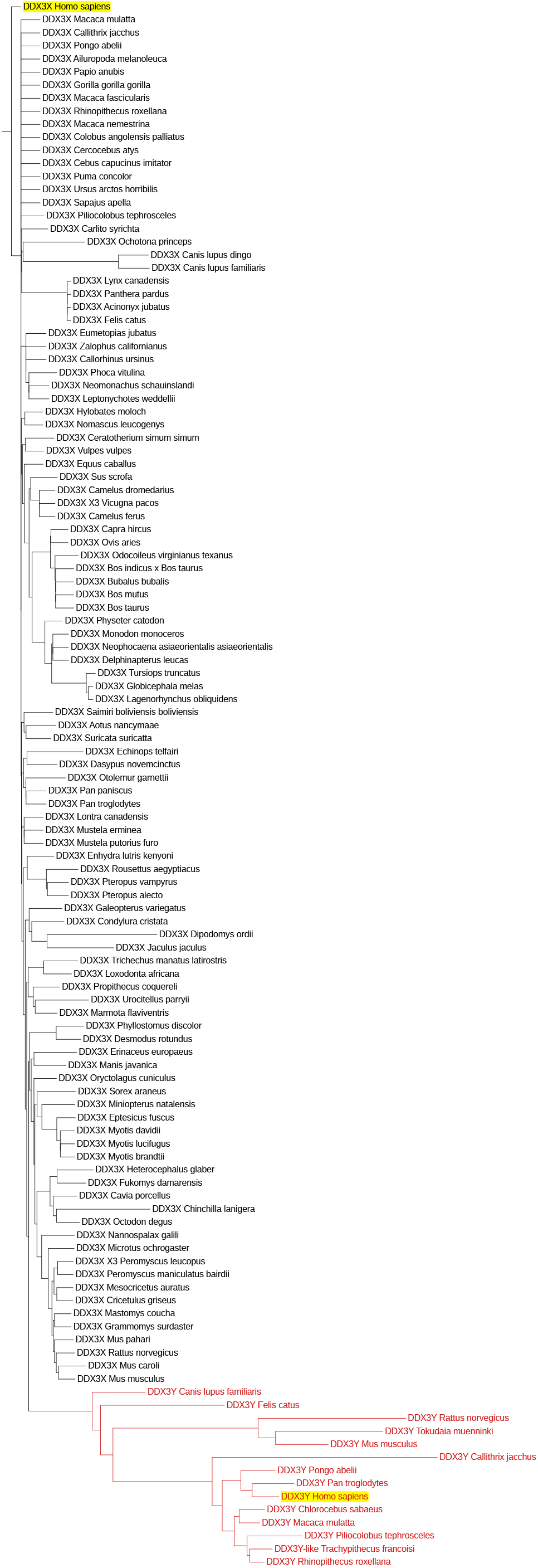
Phylogenetic tree indicating distances between the sequences of DDX3X and DDX3Y in mammals (where sequence is available). Human DDX3X and DDX3Y are highlighted in yellow. The cluster of mammalian DDX3Y orthologs is indicated by red branches.

**Figure S2:**
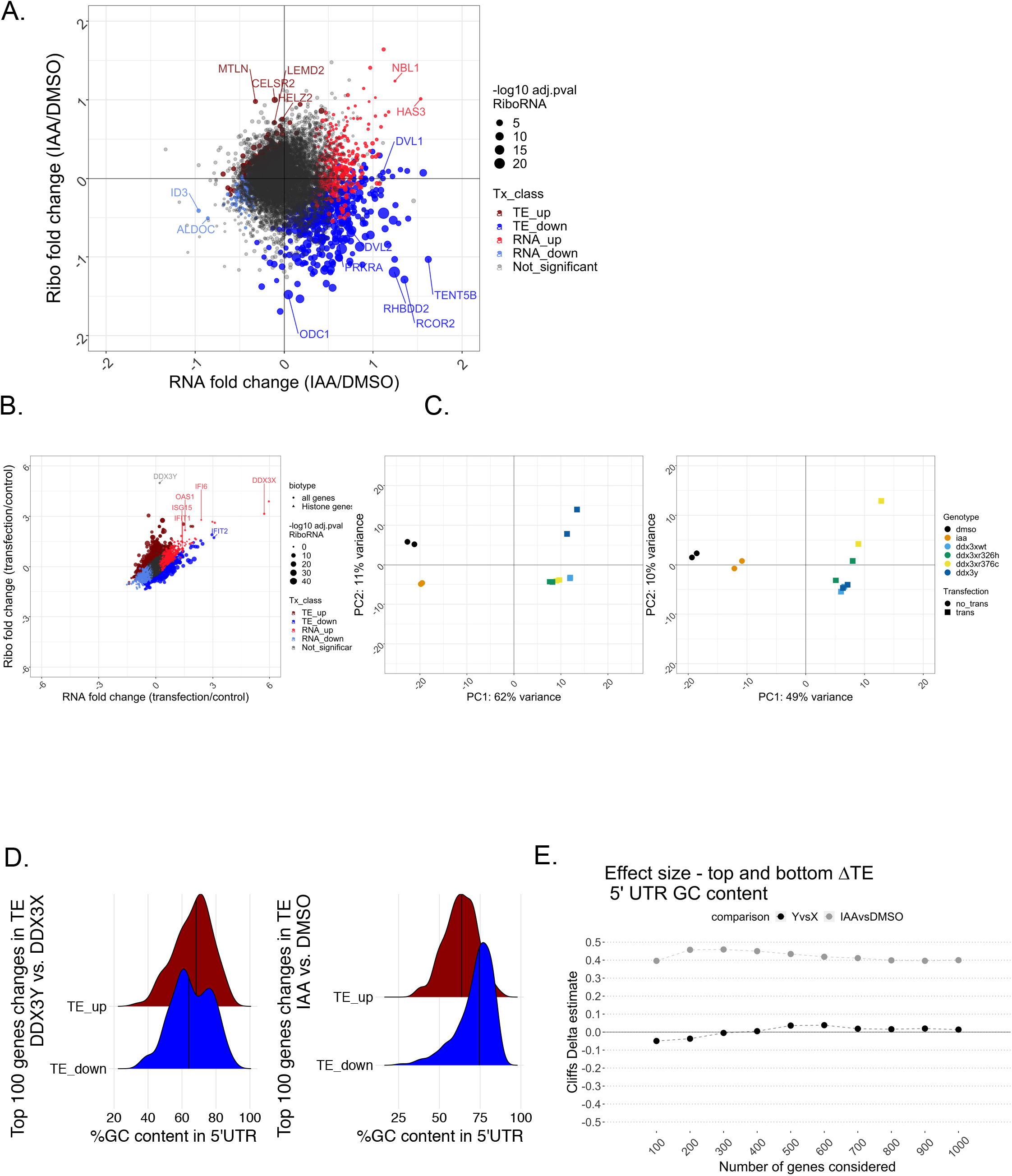
**A**. Differential expression analysis of RNA and ribosome profiling changes upon DDX3 degradation. Point size indicates differential translation **B**. Effects of transfection on ribosome occupancy and RNA levels in HCT116 cells. DDX3 paralogs and selected interferon response related genes are labelled. **C**. Principal component plots of **Left:** RNA levels and **Right:** ribosome occupancy in HCT116 cells transfected with DDX3 variants. The major source of variation is the transfection itself. **D**. GC content in the 5’ UTR of the 100 genes with the greatest magnitude of increase (TE_up) or decrease (TE_down) in translation efficiency upon expression of DDX3Y vs. DDX3X or IAA vs. DMSO (without accounting for statistical significance). **E**. Effect size (Cliff’s delta) between 5’UTR GC content of gene sets with indicated Ns selected as in (A). Number of genes indicated on the X-axis.

**Figure S3:**
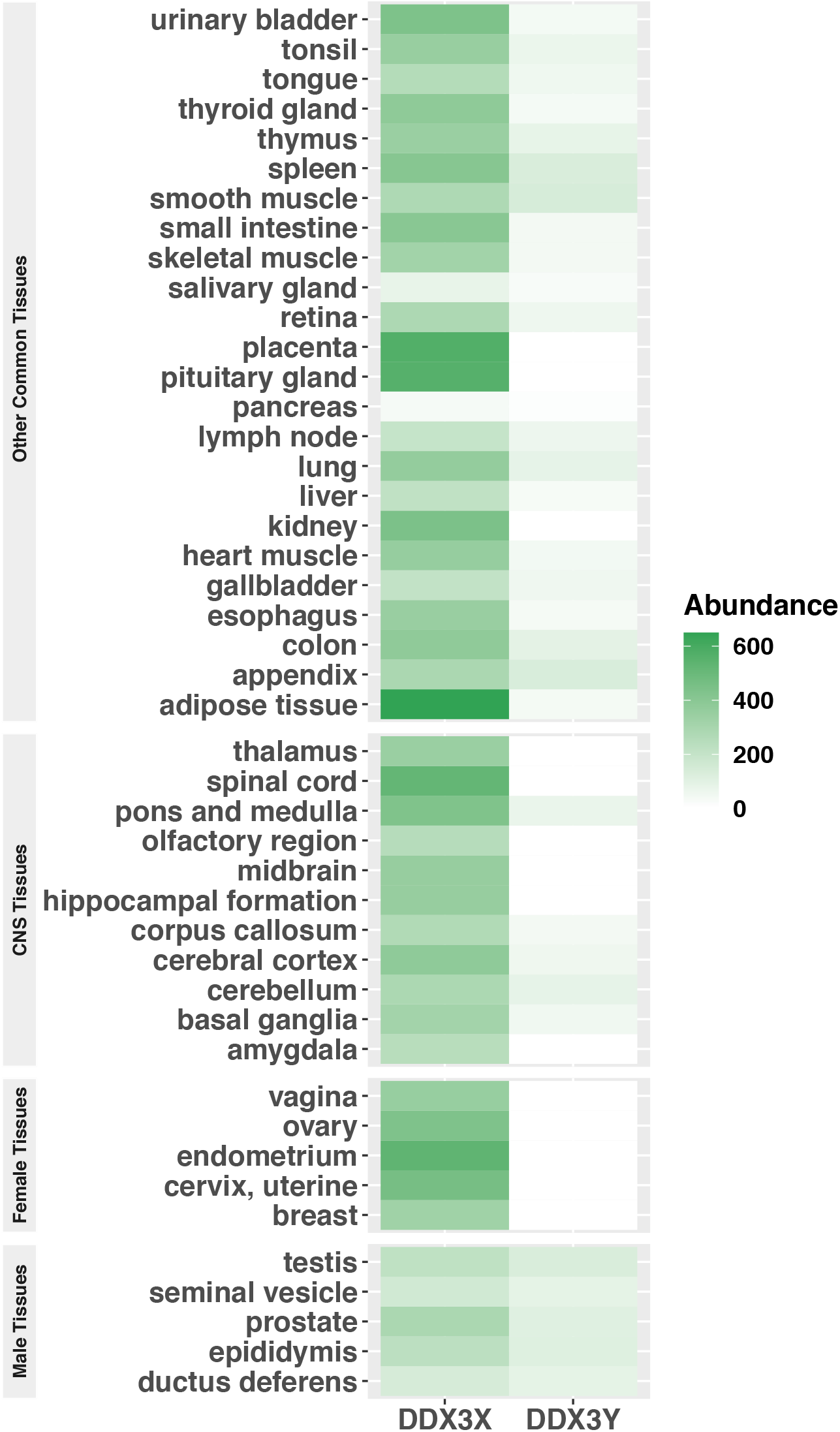
Scaled TPM expression of DDX3X and DDX3Y across a number of human tissues (data from GTeX).

